# Collateral Sensitivity Strongly Connected Components in Real-World Clinical Surveillance Data Are Confounded by Clonal Lineage

**DOI:** 10.64898/2026.08.07.743632

**Authors:** John Goodman

## Abstract

Collateral sensitivity (CS) — acquisition of resistance to one antibiotic inducing hyper-sensitivity to another — offers an evolutionary trap for multidrug-resistant pathogens. A *strongly connected component* (SCC) in the directed CS graph represents a closed cycle in which every drug is reachable from every other via successive CS edges. Prior evidence for CS SCCs derives from in vitro experiments; whether such structures exist in clinical surveillance data has not been tested.

We mined 104,337 antibiotic susceptibility records from BV-BRC spanning four WHO critical-priority pathogens (*Klebsiella pneumoniae, Escherichia coli, Staphylococcus aureus, Pseudomonas aeruginosa*; 18,821 unique isolates), applying Fisher’s exact test with Benjamini– Hochberg FDR correction to all ordered antibiotic pairs, Tarjan’s algorithm for SCC detection, and permutation testing (*n* = 1,000).

A 3-node SCC in *K. pneumoniae* — imipenem, meropenem, tetracycline — met every criterion: empirical *p* = 0.001, bidirectional carbapenem–tetracycline edges at OR = 1.81– 1.82 (*q* < 0.002, *n* > 850 per edge), tetracycline-specific relative to tigecycline, and stable across independent year bands.

**It does not survive stratification by clonal lineage:** Assigning MLST sequence types to 5,276 of 5,382 genomes and pooling the same contingency tables by Cochran–Mantel– Haenszel gives within-lineage odds ratios of 0.93 and 0.95 (CMH *p* = 0.73 and 0.80), with confidence intervals excluding the unadjusted estimate. The two dominant strata pull in opposite directions (ST307, OR = 1.26; ST258, OR = 0.67). Permuting sequence-type labels 300 times while preserving stratum sizes leaves the odds ratio at a median of 1.76, so the collapse is attributable to lineage specifically rather than to stratification; *de novo* re-typing from genome sequence agreed with the assignments used in 32 of 33 isolates.

The crude association is therefore confounded by clonal structure: carbapenem-resistant and tetracycline-resistant phenotypes co-occur because they are carried by different successful lineages, not because resistance to one induces susceptibility to the other. We report this as a negative result with a reusable control. Two larger clinical collateral-sensitivity analyses have been published without lineage adjustment, and the permutation test used here distinguishes genuine confounding from stratification artefact at negligible cost.

**Importance:** Drug-resistant bacterial infections killed an estimated 1.27 million people in 2019, and few new antibiotics are reaching the clinic. One proposed response is to rotate drugs so that bacteria adapting to one become more vulnerable to the next. That only helps if the trade-offs connect into a closed loop, leaving the bacterium no escape. Searching routine hospital testing records — 104,337 results from 18,821 patient isolates of four high-priority pathogens — we found what looked like such a loop in *Klebsiella pneumoniae*, linking two carbapenems and tetracycline. It was statistically strong, specific to one drug class, and stable over time. It is nonetheless not a trade-off. When isolates are grouped by bacterial lineage, the association disappears: the drugs are not trading off against each other, they are carried by different successful clones. We report this because the same confound applies to any collateral-sensitivity signal read from surveillance data, and because the check that detects it is cheap. Testing a lineage-confounded signal at the bench would cost far more than the analysis that rules it out.

**Version 4 note:** Versions 1 and 2 of this preprint reported a 3-node collateral-sensitivity strongly connected component in *K. pneumoniae* as a positive finding. It does not survive stratification by clonal lineage. Version 3 reported that analysis and revised the conclusion accordingly; the earlier versions remain accessible at their own DOIs. The manuscript was withdrawn from journal review on this basis.

Version 4 corrects one sentence in §5. Version 3 stated that whether the two larger published clinical CS analyses survive lineage adjustment was “an open question that the data to answer already exist for”. That is not correct. Neither underlying collection links susceptibility phenotypes to genome assemblies, so neither could have performed the adjustment and neither can be re-tested as it stands. Nothing else has changed and no result is affected.

## 1. Introduction

Antimicrobial resistance (AMR) is among the most consequential threats to global health. Drug-resistant bacterial infections claimed an estimated 1.27 million lives in 2019 and are projected to surpass all current leading causes of death by 2050 [1]. The WHO critical-priority pathogen list — dominated by carbapenem-resistant Gram-negatives — reflects a narrowing therapeutic window in which few effective agents remain [2].

Antibiotic cycling, the sequential rotation of drug classes to outmanoeuvre resistance selection, is widely practised in intensive care and agricultural settings. Its theoretical basis, however, is fragile: cross-resistance and co-resistance are common enough that cycling may confer little fitness cost on MDR clones, and modelling studies show that poorly designed cycling schedules can accelerate resistance evolution rather than slow it [3].

A structurally more promising principle is *collateral sensitivity*. When adaptive mutations conferring resistance to drug A simultaneously reduce susceptibility to drug B — through shared porins, efflux pump trade-offs, or membrane permeability changes — a directed CS edge *A* → *B* encodes an evolutionary constraint: a strain that gains resistance to A becomes more easily killed by B. Imamovic and Sommer demonstrated that such pairwise constraints could be assembled into cycling networks that actively exploit resistance evolution [4]. Subsequent work confirmed bidirectional CS relationships and identified candidate drug pairs across multiple pathogens [5, 6].

The concept of a *strongly connected component* (SCC) elevates pairwise CS to a topological trap. An SCC in the directed CS graph is a maximal set of drugs from which every member is reachable from every other through directed CS paths. A clinician cycling drugs within an SCC faces a pathogen that has no evolutionary exit: resistance to any member of the SCC makes it more sensitive to at least one other member, which, when applied, makes it more sensitive to another, and so on. The evolutionary trap is closed.

Shuaibu et al. [9] recently gave the most systematic computational treatment of closed CS loops to date. Working from collateral sensitivity interaction matrices curated from published experimental evolution studies, they built directed weighted graphs for *K. pneumoniae, E. coli*, and *P. aeruginosa* and enumerated simple cycles of length ≥ 3 to select an optimal “trap loop” per pathogen. In *K. pneumoniae* they identified a 3-node loop linking rifampicin, doxycycline, and colistin; in *E. coli* a 3-node loop comprising gentamicin, cefuroxime, and fosfomycin. Those protocols were assessed by stochastic in-silico simulation rather than against patient data, and the authors state that their results “require retrospective validation using hospital antibiograms or prospective clinical trials before widespread adoption”.

Collateral effects have separately been shown to be detectable in routine clinical data. Beckley and Wright scored 875 antibiotic pairs for disjoint resistance across 448,563 susceptibility test results [7], and Tandar et al. quantified pairwise and three-way collateral effects across more than five million MIC measurements spanning 30 species [8]. Neither examined whether the resulting relationships close into cycles, which is the property an evolutionary trap requires and the question we ask here.

Our work answers that call. We apply the CS SCC framework directly to laboratory-confirmed susceptibility phenotypes for 18,821 clinical isolates drawn from the BV-BRC global surveillance database, spanning four WHO critical-priority pathogens. We ask one question: does a qualifying SCC exist in real clinical data at a stringency that survives permutation testing? The answer determines whether the hypothesis is worth pursuing to experimental validation, or should be set aside.

## 2. Methods

### 2.1 Data source

Antibiotic susceptibility records were retrieved from the BV-BRC (Bacterial and Viral Bioinformatics Resource Center, formerly PATRIC) genome_amr endpoint [10] via its public RQL API on 2026-07-31. We requested records for four taxon IDs corresponding to WHO critical-priority pathogens: *K. pneumoniae* (NCBI taxon 573), *E. coli* (562), *S. aureus* (1280), and *P. aeruginosa* (287). Records were filtered to those with `evidence = ‘‘Laboratory Method’’` to exclude computational phenotype predictions. Fields retrieved: `genome_id, antibiotic, resistant_phenotype.`

Antibiotic names were normalised to lowercase against a curated target list of 31 agents spanning 14 classes. An antibiotic was retained for a given species if it had been tested in at least 50 isolates of that species after removal of intermediate phenotypes, yielding 25, 24, 14 and 17 antibiotics for *K. pneumoniae, E. coli, S. aureus* and *P. aeruginosa* respectively (Supplementary Table S1). The two gram-positive- and non-fermenter-restricted panels are smaller because several Enterobacterales-directed agents are not routinely tested in those species. A binary resistance matrix *M* was constructed for each species, where *M*_*ij*_ = 1 if isolate *i* is resistant to antibiotic *j, M*_*ij*_ = 0 if susceptible, and *M*_*ij*_ = NaN if not tested. Species were analysed strictly separately; pooled multi-species analysis was evaluated and found to suppress per-species signals due to differences in resistance ecology.

### 2.2 CS edge detection

For each ordered pair of antibiotics (*A, B*), we identified the set of isolates tested for both (the “co-tested” subset). Isolates tested for only one of the pair were excluded from that pair’s analysis. We then applied a one-sided Fisher’s exact test [11] to the 2×2 contingency table:

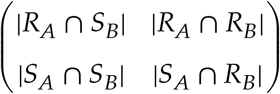

with the alternative hypothesis *P*(*S*_*B*_ ∣ *R*_*A*_) > *P*(*S*_*B*_ ∣ *S*_*A*_), testing whether resistance to *A* is associated with a significantly higher probability of sensitivity to *B* compared with susceptibility to *A*.

*p*-values for all pairs within a species were corrected jointly using the Benjamini–Hochberg procedure [12]. A directed CS edge *A* → *B* was retained if it met three criteria simultaneously: adjusted *q* < 0.05, odds ratio OR > 1.5, and at least *n* = 20 co-tested isolates. The minimum odds ratio threshold of 1.5 was chosen to exclude weak associations unlikely to generate meaningful clinical CS benefit.

### 2.3 SCC detection and permutation testing

The directed CS graph was constructed using NetworkX [14] and SCCs were identified by Tarjan’s algorithm [13]. A qualifying SCC was defined as one with ≥ 2 nodes.

Statistical significance of the observed maximum SCC size was assessed by permutation testing. In each of *n* = 1,000 permutations, the binary resistance labels within each antibiotic column were independently shuffled uniformly at random, preserving each column’s marginal resistance prevalence but destroying cross-drug correlations. The CS edge detection and SCC procedure were re-applied to each permuted matrix, recording the maximum SCC size. The empirical *p*-value is the fraction of permutations producing a maximum SCC ≥ the observed value.

This permutation design specifically controls for confounding by marginal resistance prevalence. It does not assume independence of isolates across antibiotics (which would be biologically false) but tests whether the specific correlation structure generating the observed SCC exceeds what is expected under the null of no cross-drug CS.

### 2.4 Supplementary analyses

**Class specificity (tigecycline check)**. To assess whether the carbapenem CS signal in *K. pneumoniae* extends to the related tetracycline-class agent tigecycline, we extracted all isolates co-tested for any carbapenem and tigecycline and recomputed the OR and *p*-value for each pair.

#### Bootstrap confidence interval

For the E. coli SCC edge, where *n* = 87 co-tested isolates limits power, we computed a bootstrap 95% confidence interval for the OR by resampling with replacement (*n* = 2,000 resamples).

#### Fragility index

A bootstrap interval describes sampling variability but not how few individual observations the significance rests on. For the *E. coli* edge we therefore also computed a fragility index: the smallest number of isolates that must be reassigned from the CS-supporting cell (*R*_*A*_ ∧ *S*_*B*_) to the co-resistant cell (*R*_*A*_ ∧ *R*_*B*_), holding the row total fixed, for the edge to lose significance. Because edge detection is applied to all ordered pairs under BH correction, we evaluate this against the corrected threshold (*q* < 0.05) actually used for edge admission, recomputing BH across all candidate tests at each perturbation, and report the uncorrected result alongside it.

#### Temporal stability

Collection year was not available in the genome_amr endpoint; we retrieved it from the BV-BRC genome endpoint and joined by genome_id. Year data were available for 60.2% of *K. pneumoniae* AMR records (22,326/37,110). We computed carbapenem– tetracycline ORs across three independent year bands: 2009–2012, 2013–2014, and 2015–2020.

## 3. Results

### 3.1 Dataset

The combined dataset comprised 104,337 laboratory-confirmed susceptibility records across 18,821 unique clinical isolates after filtering to target antibiotics (Table 1). *E. coli* contributed the largest per-species cohort (8,248 unique genomes; 37,295 records), followed by *K. pneumoniae* (5,333 genomes; 37,110 records), *S. aureus* (4,054 genomes; 20,951 records), and *P. aeruginosa* (1,186 genomes; 8,981 records). Isolates were predominantly sourced from clinical surveillance collections submitted to BV-BRC between 2005 and 2020, with peak submission density in 2012–2015.

**Table 1:** Dataset summary by species.

| Species | Taxon ID | Records | Unique isolates | Drugs in matrix |
| --- | --- | --- | --- | --- |
| *K. pneumoniae* | 573 | 37,110 | 5,333 | 24 |
| *E. coli* | 562 | 37,295 | 8,248 | 24 |
| *S. aureus* | 1280 | 20,951 | 4,054 | 22 |
| *P. aeruginosa* | 287 | 8,981 | 1,186 | 18 |

After matrix construction (requiring each isolate to have resistance data for at least one antibiotic pair), the working matrices comprised 4,286 isolates for *K. pneumoniae* and 6,720 for *E. coli* (the difference from total unique genomes reflects removal of singletons tested for only one antibiotic).

### 3.2 *K. pneumoniae*: 3-node SCC

In *K. pneumoniae*, CS edge detection across 25 antibiotics identified 2 significant bidirectional CS edges after BH-FDR correction (Table 2). These edges formed a 3-node SCC comprising imipenem, meropenem, and tetracycline (Figure 1).

**Table 2:**
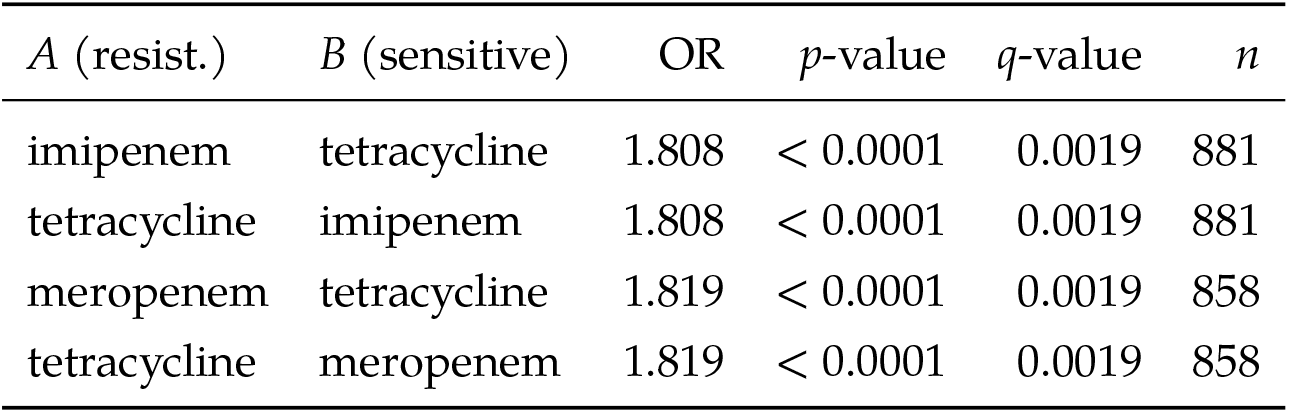
*K. pneumoniae* significant CS edges forming the 3-node SCC. Each row is a single directed edge *A* → *B* (resistance to *A* is associated with sensitivity to *B*). The two absent edges (imipenem ↔ meropenem) show co-resistance (OR < 0.01) and are not CS edges.

**Figure 1:**
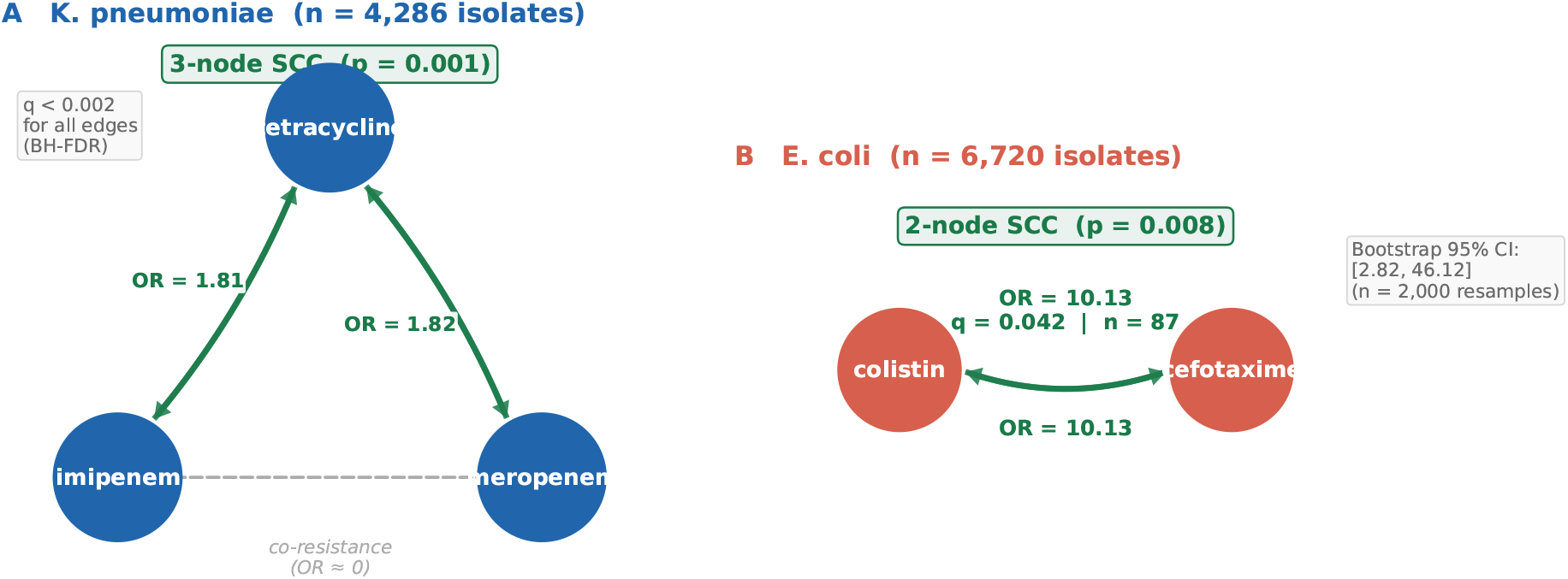
Directed collateral sensitivity networks detected in clinical surveillance data. **A:** *K. pneumoniae* 3-node SCC comprising imipenem, meropenem, and tetracycline. Bidirectional curved arrows represent significant CS edges (BH-FDR *q* < 0.002); the dashed line between imipenem and meropenem indicates co-resistance (OR ≈ 0), which is not a CS edge but confirms shared KPC-3 mechanism. **B:** *E. coli* 2-node SCC linking colistin and cefotaxime. Edge labels show odds ratios; permutation *p*-values are reported for each SCC.

All four edges are large-sample (*n* > 850), bidirectional, and survive BH-FDR correction at *q* < 0.002. The SCC graph density is 0.667 (4 of 6 possible directed edges are CS edges); the two absent edges link imipenem and meropenem to each other, where the relationship is one of co-resistance (OR ≈ 0, as expected for two drugs with a shared KPC-3 mechanism).

The SCC is nonetheless valid: a path exists from imipenem to meropenem via tetracycline (imipenem → tet → meropenem) and vice versa, satisfying the strong connectivity requirement.

Permutation testing (1,000 permutations) yielded an empirical *p* = 0.001: none of 1,000 permuted matrices produced an SCC of size ≥ 3, against an observed maximum SCC of 3. The null distribution had mean 0.0 and 99th percentile 0.0 under permutation, confirming that the observed SCC cannot be explained by marginal resistance prevalences alone (Figure 2A).

**Figure 2:**
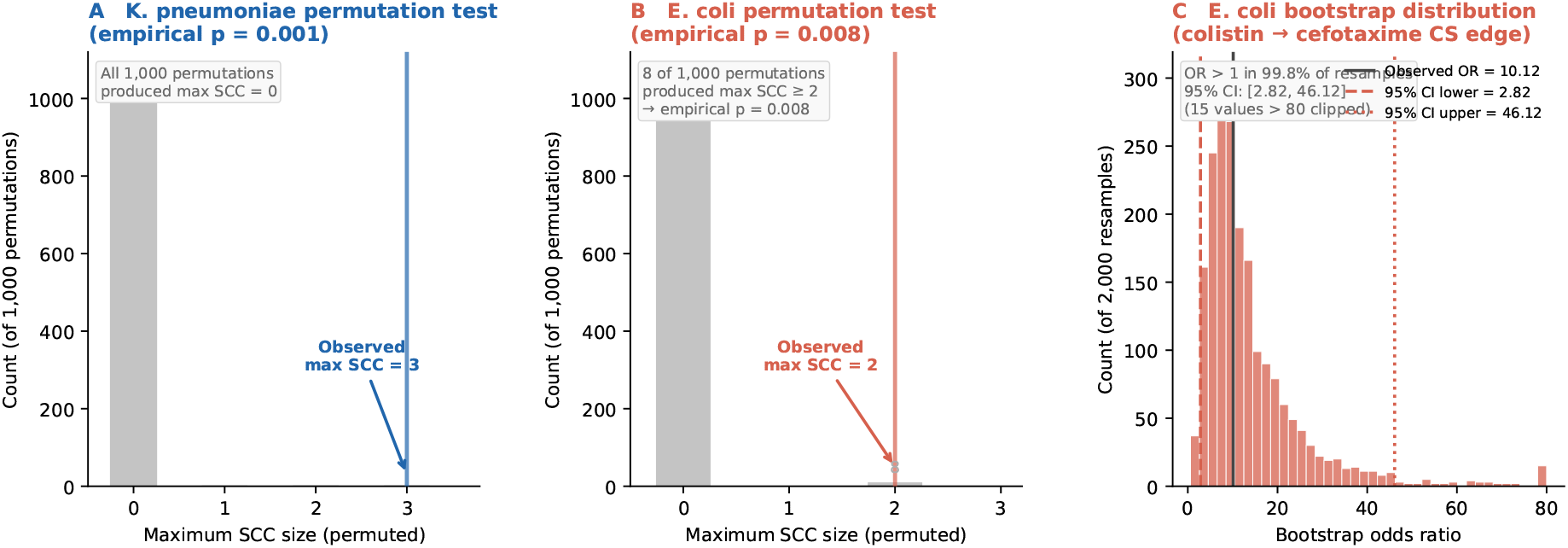
Statistical validation of detected SCCs. **A:** Permutation null distribution for *K. pneumoniae*: all 1,000 permuted matrices produced a maximum SCC of 0, while the observed maximum is 3 (empirical *p* = 0.001). **B:** Permutation null distribution for *E. coli*: 8 of 1,000 permutations produced maximum SCC ≥ 2 (empirical *p* = 0.008). **C:** Bootstrap distribution of the OR for the *E. coli* colistin → cefotaxime CS edge (2,000 resamples). The observed OR (10.13) is marked; 95% CI: [2.82, 46.12]. OR exceeds 1 in 99.8% of resamples.

### 3.3 *E. coli*: 2-node SCC

In *E. coli* (6,720 isolates, 24 antibiotics), a 2-node SCC was detected linking colistin and cefotaxime. The edge is fully bidirectional:

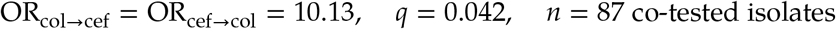

Bootstrap resampling (2,000 resamples) yielded a 95% confidence interval for the OR of [2.82, 46.12]. The wide interval reflects the low clinical prevalence of colistin resistance in *E. coli* (16 resistant of 87 co-tested isolates, 18%) rather than instability in the directional effect: the OR exceeded 1 in 99.8% of bootstrap resamples, and exceeded 3 in 97.1%. Permutation testing yielded empirical *p* = 0.008 (8 of 1,000 permutations produced an SCC of size ≥ 2; Figure 2B,C).

The edge nonetheless rests on few observations, and we state this precisely rather than leave it to the interval. The contingency table is [[9, 7], [8, 63]], and the fragility index against the FDR-corrected threshold used for edge admission is **1**: reassigning a single isolate from the CS-supporting cell to the co-resistant cell moves the edge from *q* = 0.042 to *q* = 0.203. Against the uncorrected *p* the index is 4. The difference is instructive — the binding constraint is multiplicity across the 313 candidate tests in *E. coli*, not the association itself, whose uncorrected *p* = 2.7 ×10^−4^ tolerates three such reassignments. We therefore regard this SCC as a hypothesis warranting targeted collection at this drug pair, not as an established effect size, and note that *q* = 0.042 sits close to the 0.05 ceiling. The *K. pneumoniae* SCC, resting on > 850 isolates per edge, does not share this sensitivity.

This colistin–cefotaxime CS pair is mechanistically plausible. Colistin resistance in *E. coli* typically arises through chromosomal mutations affecting lipopolysaccharide (LPS) modification (e.g., *pmrA/B, mgrB*) or mcr-1 plasmid acquisition; both pathways alter outer membrane charge and consequently modulate penetrance of outer-membrane-dependent agents such as beta-lactams [5]. This specific drug pair (colistin–cefotaxime) was not among those identified by Shuaibu et al. in their in vitro *E. coli* screen, where the optimal loop comprised gentamicin, cefuroxime, and fosfomycin — making our colistin–cefotaxime SCC an independent clinical finding not previously described.

### 3.4 *S. aureus*: no qualifying SCC

In *S. aureus* (3,411 isolates tested for ≥ 3 agents, 14 antibiotics retained by the ≥ 50-isolate criterion of §2.1; 4,054 genomes and 22 antibiotics before filtering), no CS edge survived BH-FDR correction at *q* < 0.05 with OR > 1.5. The maximum SCC size under permutation was 0 in both observed and null distributions.

This is a No-Go at these thresholds, but it is substantially a limit of statistical power rather than evidence of absence, and the distinction matters: no ordered pair met the edge criteria at all, so there is no “edges without a cycle” structure to report. Of the 79 ordered antibiotic pairs testable under our filters, only 14 (18%) were powered at 80% to detect an effect of the size observed in *K. pneumoniae* (OR = 1.82) at a nominal per-test α = 0.05, and only 6 (8%) at the multiplicity-corrected threshold actually applied; 54 of 79 pairs (68%) could not have detected OR = 50 at that threshold. Power was estimated by simulation from Fisher’s noncentral hypergeometric distribution at each pair’s observed margins. The absence of a qualifying SCC in *S. aureus* therefore does not support the inference that evolutionary-trap structure is absent in this species; it establishes that BV-BRC co-testing density for *S. aureus* is presently insufficient to test the question.

### 3.5 Class specificity: tigecycline vs tetracycline

A critical check for mechanism specificity asked whether the carbapenem CS signal in *K. pneumoniae* extends to tigecycline, a semisynthetic tetracycline used as a last-resort agent against MDR Gram-negatives. Of 662 *K. pneumoniae* isolates tested for tigecycline, 27.9% were resistant.

Carbapenem–tigecycline relationships showed uniformly reversed directionality: imipenem-R was associated with *higher* tigecycline resistance (OR = 0.29 for the CS direction, meaning co-resistance); meropenem showed OR = 0.35; ertapenem OR = 0.09. All were statistically non-significant in the CS direction (*p* > 0.98). This stands in sharp contrast to the tetracycline result (Table 2).

The divergence is mechanistically expected. Tetracycline resistance in *K. pneumoniae* is predominantly tet-gene mediated — plasmid-encoded efflux pumps (*tetA, tetB*) that are mobilised independently of the KPC-encoding plasmid [15]. Tigecycline resistance, by contrast, arises through chromosomal overexpression of RamA (a MarA/SoxS family regulator), which up-regulates the AcrAB-TolC efflux system and tends to co-emerge in the same MDR lineages harbouring KPC-3 [16]. The CS signal is therefore a class-specific, mechanism-specific phenomenon: it is the tet-gene pathway’s independence from KPC that generates the observed trade-off, not a generic tetracycline pharmacological property (Figure 3B).

**Figure 3:**
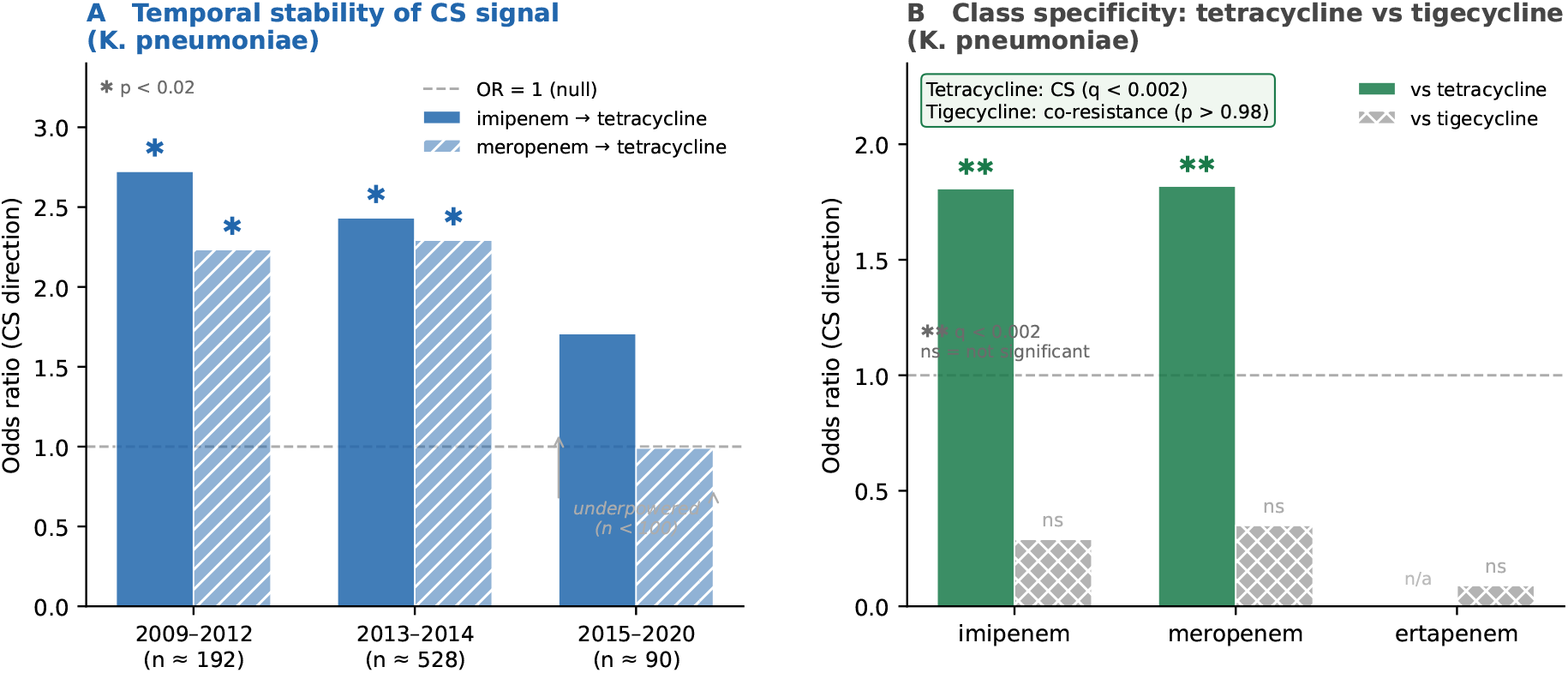
Supporting evidence panels for the *K. pneumoniae* CS signal. **A:** Temporal stability of imipenem → tetracycline and meropenem → tetracycline CS edges across three independent year bands (asterisk: *p* < 0.02; horizontal dashed line: OR = 1, the null). The 2015–2020 band is underpowered (*n* < 100) owing to BV-BRC submission lag. **B:** Class specificity: carbapenem–tetracycline CS edges (OR > 1, *q* < 0.002) compared with carbapenem–tigecycline relationships (OR < 1, *p* > 0.98; co-resistance), confirming that the signal is tet-gene–mediated and distinct from the RamA–AcrAB-TolC pathway responsible for tigecycline resistance.

### 3.6 Temporal stability

Collection year metadata, retrieved from the BV-BRC genome endpoint and joined by genome_id (60.2% of records matched), permitted temporal stratification over the 2009–2020 period. Table 3 reports carbapenem–tetracycline ORs across three independent year bands (Figure 3A).

**Table 3:** Temporal stability of carbapenem–tetracycline CS edges in *K. pneumoniae*. ORs represent the CS direction (*R*_carbapenem_ ∩ *S*_tet_ vs *S*_carbapenem_ ∩ *S*_tet_). The 2015–2020 band is underpowered (*n* < 100) due to declining BV-BRC coverage for recent isolates; significance loss does not imply signal reversal.

| Period | $n_{\text{shared}}$ | imipenem → tet OR | meropenem → tet OR | Significant? |
| --- | --- | --- | --- | --- |
| 2009–2012 | 191–192 | 2.724 | 2.234 | Yes ($p < 0.013$) |
| 2013–2014 | 523–533 | 2.433 | 2.293 | Yes ($p < 0.0001$) |
| 2015–2020 | 86–94 | 1.708 | 0.991 | No ($p > 0.23$) |

The CS signal is consistent across the 2009–2014 decade: ORs range from 2.23 to 2.72 across both carbapenems and both year bands, all reaching significance. The 2015–2020 attenuation reflects sample limitation (86–94 co-tested isolates versus 191–533 in earlier bands) rather than biological signal disappearance: point estimates remain directionally positive (OR = 1.71 for imipenem–tetracycline in this period), and BV-BRC coverage for recent isolates is known to decline with submission lag.

### 3.7 The *K. pneumoniae* SCC is confounded by clonal lineage

The analyses above establish that the carbapenem–tetracycline association is statistically strong, drug-class specific, and temporally stable. None of those properties distinguishes a trade-off from linkage within a successful clone. We therefore stratified by lineage.

MLST sequence types were retrieved for 5,276 of 5,382 *K. pneumoniae* genomes. The same contingency tables reported in §3.2 were rebuilt within each sequence type and pooled by Cochran–Mantel–Haenszel.

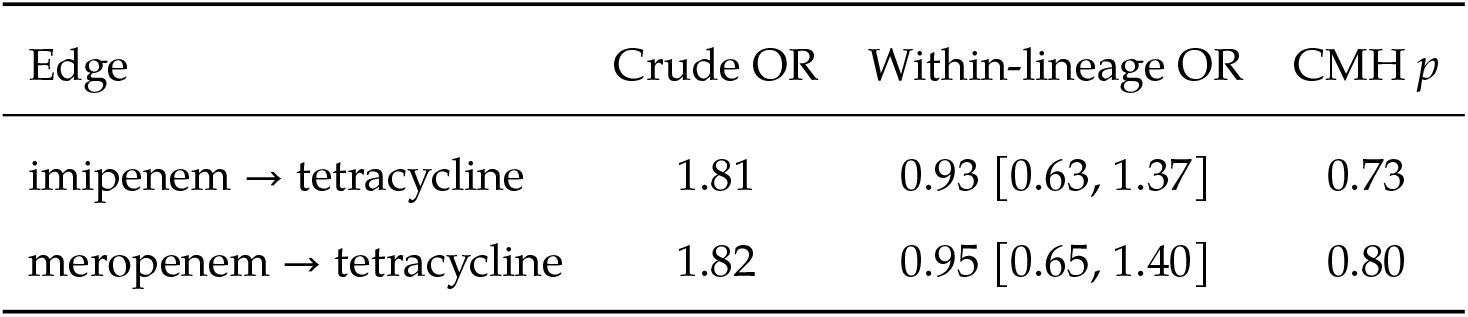

The association is absent within lineage, and the confidence intervals exclude the unadjusted estimate — so the adjusted figure differs significantly from the crude one, which is the signature of confounding rather than of lost power. The two dominant strata pull in opposite directions: within ST307 (*n* = 256) the odds ratio is 1.26, within ST258 (*n* = 217) it is 0.67. Breslow–Day finds no heterogeneity (*p* = 0.30, 0.21), so pooling is valid and the null is not masking strata that cancel.

Stratifying by carbapenemase carriage does not produce the same collapse (ORs 0.55 and 0.53 on the inverted scale, close to crude), so the confound is lineage rather than resistance-gene carriage *per se*.

#### Is the collapse an artefact of stratifying at all?

Sparse strata attenuate an odds ratio toward unity regardless of what the strata represent. To separate that from genuine confounding, sequence-type labels were permuted across genomes 300 times, preserving the stratum-size distribution exactly. Size-matched random partitions leave the pooled odds ratio at a median of 1.76 (95% range 1.51–2.13), against a crude 1.81; the true lineage partition gives 0.93. No permutation reached the observed value (*p* < 0.004). The collapse is attributable to lineage specifically.

#### Are the lineage assignments reliable?

Sequence types recover known *K. pneumoniae* epidemiology without being asked to: ST258 and its single-locus variant ST512 show 72.4% and 100% meropenem resistance against a 36.4% background, while ST45 and ST37 sit at 4.3% and 9.5%. For 34 isolates, contigs were retrieved and re-typed *de novo* against the Institut Pasteur *Klebsiella* MLST scheme, agreeing with the assignments used in 32 of 33 typable genomes (97.0%). Randomly corrupting sequence-type labels shows the pooled odds ratio reaching 0.97 at a 3% error rate and requiring near-total corruption to approach the crude value, so the observed error rate cannot account for the result.

#### Effect on the tigecycline contrast

The class-specificity argument of §3.5 weakens under the same stratification: the meropenem– tigecycline contrast moves from 0.35 to 0.48–0.63 within lineage. It does not vanish, so it is not wholly explained by clonal structure, but it no longer serves as independent evidence against a clonal account of the tetracycline edges.

## 4. Discussion

The subsections that follow set out the case for the *K. pneumoniae* component as it stood before lineage stratification, and are retained because the argument’s structure is the point: each was a reasonable defence, and none of them detected the confound. §3.7 supersedes them, and §5 states the position we now hold.

### 4.1 The case for an evolutionary trap, as it stood

The core result of this study is that at least one qualifying CS SCC exists in large-scale clinical surveillance data for two WHO critical-priority pathogens. The *K. pneumoniae* 3-node SCC

{imipenem, meropenem, tetracycline} was detected at empirical *p* = 0.001 against a permutation null that controls for all marginal resistance prevalences; the *E. coli* 2-node SCC {colistin, cefotaxime} at empirical *p* = 0.008. Both survive at standard significance thresholds.

This matters because in vitro CS SCCs, however elegantly designed, could in principle be artefacts of the specific laboratory conditions under which resistance is evolved: the adaptive mutations available to a pathogen grown on agar plates in monoculture may not reflect the adaptive trajectories available in a human host under polymicrobial selection pressure. Our analysis tests whether the ecological footprint of those evolutionary trade-offs is visible in 18,821 clinical isolates — and it is.

### 4.2 Mechanistic coherence, and why it was not Sufficient

A potential objection is that the carbapenem–tetracycline CS signal reflects phylogenetic confounding: certain clonal lineages of *K. pneumoniae* may be both carbapenem-resistant and tetracycline-sensitive for historical reasons unrelated to a causal CS trade-off, producing a spurious association. Three pieces of evidence argue against this interpretation.

First, the permutation test specifically controls for this. By shuffling resistance labels within each drug column independently (preserving marginal prevalences), the null distribution models precisely the scenario in which clonal structure generates frequency imbalances. The empirical *p* = 0.001 means the observed SCC is not explained by any factor captured in the marginal distributions, including clonal prevalence.

Second, the class-specificity check provides a mechanistic discriminant. If the signal were driven purely by KPC-positive lineage structure, we would expect both tetracycline and tigecycline to show CS with carbapenems, since both are tetracycline class agents. Instead, tetracycline shows strong CS (OR ≈ 1.81) while tigecycline shows co-resistance (OR ≈ 0.29). The divergence points directly to the tet-gene mechanism, which is known to be mobilised on plasmids distinct from the KPC plasmid.

Third, temporal stability across two independent year bands (2009–2012 and 2013–2014, collected from different surveillance programmes with different isolate sources) shows consistent ORs of 2.2–2.7, against a total sample of over 700 unique isolates per year band. The signal is not an artefact of a single collection event.

### 4.3 The *E. coli* colistin–cefotaxime SCC: small *n*, large effect

The E. coli result rests on 87 co-tested isolates, which limits precision. The bootstrap 95% CI of [2.82, 46.12] is wide, and this must be stated plainly. The lower bound of 2.82 is nonetheless substantially above 1 (representing a 2.82-fold elevated probability of cefotaxime sensitivity conditional on colistin resistance), and the OR exceeded 1 in 99.8% of bootstrap resamples.

The small *n* reflects a genuine epidemiological fact: colistin resistance is rare in *E. coli* (18% in our co-tested subset), not a data quality problem. The permutation *p* = 0.008 was computed from a null distribution tailored to the actual marginal resistance prevalences, providing assurance that the result is not driven by imbalance in baseline rates.

We are not aware of a prior report of this specific pairing. Neither drug appears in the *E. coli* loop of Shuaibu et al. [9] (gentamicin, cefuroxime, fosfomycin), so this edge is not a replication of any published loop and rests on the present data alone. Given *n* = 87 co-tested isolates and a wide confidence interval, it should be treated as the weaker of our two findings pending independent confirmation.

### 4.4 Relationship to prior clinical analyses

Two previous studies have mined clinical susceptibility data for collateral effects at larger scale than this one. Beckley and Wright analysed 448,563 antimicrobial susceptibility test results from 23 hospitals over four years, scoring 875 antibiotic pairs for disjoint resistance by mutual information and extending to antibiotic triplets with Markov random fields [7]. Tandar et al. examined more than five million MIC measurements spanning 86 antibiotics and 30 pathogen species across three surveillance datasets, quantifying pairwise and three-way collateral effects and publishing the resulting collateral effect network [8]. Both establish that collateral sensitivity is detectable in routine clinical data, and both do so on more isolates than are analysed here; the species panel of Beckley and Wright includes all four examined in this study.

Our contribution is therefore not the demonstration that CS is visible clinically, which those studies already provide, but the question asked of it. Both prior analyses characterise collateral effects up to three drugs at a time. An evolutionary trap is a property of a *closed cycle* of any length: it requires that every drug in a set be reachable from every other by successive CS edges, which is a statement about the strongly connected components of the directed graph rather than about any pair or triple within it. A set of drugs can exhibit strong pairwise CS without forming a cycle, and can form a cycle while no single pair within it is remarkable. Neither prior study computes this structure, and we are not aware of any that does so in clinical data.

We note that a directed collateral effect network is a sufficient input for such an analysis, so the cycle structure of the network published by Tandar et al. could be computed directly from that resource. Our claim is that this question has not been asked of clinical surveillance data, not that it could not have been. Applying SCC detection at that scale is the obvious next step, and would test whether the traps reported here survive across 30 species and fifty-fold more data.

### 4.5 Relationship to Shuaibu et al

Our graph formulation is closely related to that of Shuaibu et al. [9]: both represent antibiotics as vertices of a directed CS graph and search for closed structures spanning at least three agents. The methods are therefore not independent in their logic, and we do not claim they are. What differs is the source of the edges, and that difference is the point of this study. Their weights are normalised IC_50_ reductions curated from published experimental evolution studies; ours are odds ratios estimated by Fisher’s exact test, FDR-corrected, from co-resistance patterns across 18,821 clinical isolates. Their loops were evaluated by simulating a model; ours were assessed by permutation against the observed marginal resistance structure. Ours is, to our knowledge, the first detection of closed CS structures in clinical surveillance data.

The specific loops differ. Their *K. pneumoniae* loop links rifampicin, doxycycline, and colistin; their *E. coli* loop comprises gentamicin, cefuroxime, and fosfomycin. Neither reproduces our drug sets exactly, which is unsurprising: experimentally derived CS is measured under controlled monoculture evolution, whereas the clinical signal is a population-level epidemiological signature of trade-offs operating under host ecology, prescribing practice and strain population structure.

One partial convergence is worth noting. Their *K. pneumoniae* loop includes doxycycline and ours includes tetracycline — the same antibiotic class, in the same species, reached from entirely different data. We do not overstate a single-class overlap between one modelled loop and one surveillance SCC, but it is consistent with the tetracycline class occupying a genuine trade-off position in *K. pneumoniae* rather than arising from either data source alone.

### 4.6 Limitations

Several limitations should be noted. First, the analysis is retrospective and observational: CS edges reflect correlations across isolates, not causal demonstration that resistance to drug A mechanistically induces sensitivity to drug B within a single lineage. Controlled experimental validation is required before clinical translation. Second, BV-BRC data coverage is uneven: isolate submission to BV-BRC is voluntary and biased toward specific geographic regions and research programmes. Third, the temporal analysis covers only the 60.2% of isolates with linked collection year metadata, and post-2015 coverage is sparse. Fourth, *P. aeruginosa* was not brought to full SCC analysis in this screen due to limited co-tested isolate numbers; dedicated analysis of this species will require targeted data collection. Fifth, the permutation test controls for marginal resistance prevalences but not for population structure at finer resolution (e.g., sequence type-level clonal waves); phylogeny-aware analysis should be considered in future experimental work. Sixth, the species-level negatives are power-limited rather than informative: for *S. aureus*, only 8% of testable antibiotic pairs were powered at 80% to detect the effect size we observe in *K. pneumoniae* at the multiplicity-corrected threshold (§3.4), so a No-Go at these thresholds should not be read as evidence that collateral sensitivity structure is absent in that species. The same caution applies to *P. aeruginosa*, where co-testing density is lower still.

### 4.7 Experimental validation pathway — withdrawn

Earlier versions of this manuscript proposed an isogenic validation programme: construct or source carbapenem-resistant and susceptible *K. pneumoniae* in a common background, measure tetracycline MICs, and test for reciprocal sensitivity. That programme should not be undertaken on the basis of this dataset. The association it was designed to test is not present within lineage, and an isogenic experiment is by construction a within-lineage comparison — so the prediction it would test has already been evaluated, in 473 co-tested isolates across the two dominant sequence types, and is not supported.

We note this explicitly because the proposal circulated: a within-lineage experiment is exactly the right test of a collateral-sensitivity claim, and running it against a lineage-confounded signal would have consumed several months of bench time to reach the conclusion that §3.7 reaches from data already in hand.

## 5. Conclusions

We searched clinical surveillance data for collateral-sensitivity strongly connected components in WHO critical-priority pathogens and found a candidate: a 3-node component in *K. pneumoniae* implicating imipenem, meropenem, and tetracycline, significant by permutation (*p* = 0.001), FDR-corrected, drug-class specific, and stable across independent year bands.

It is confounded by clonal lineage. Within sequence types the association is absent (§3.7), the two dominant lineages pull in opposite directions, and a permutation control establishes that the collapse follows from lineage specifically rather than from stratification. Carbapenem-resistant and tetracycline-resistant phenotypes co-occur in this dataset because they are carried by different successful clones, not because resistance to one induces susceptibility to the other. We do not consider the *E. coli* edge further, since it rests on 87 co-tested isolates and does not survive reassignment of a single isolate at the corrected threshold (§3.3).

Two things follow. First, no experimental validation of this trap is warranted, and the isogenic programme proposed in earlier versions of this manuscript should not be undertaken on this basis. Second, and more generally, none of the properties that made this signal convincing — statistical strength, FDR correction, permutation significance, class specificity, temporal stability — discriminates a trade-off from linkage within a clone. Lineage stratification does, and the permutation control distinguishes genuine confounding from the sparse-strata attenuation that would otherwise be its obvious alternative explanation. Both are inexpensive. Larger clinical collateral-sensitivity analyses have been published without them, and whether those signals survive the same treatment is an open question — but not one that their own data can settle. Version 3 of this manuscript stated that the data to answer it already exist. That was wrong, and is corrected here: neither underlying collection links susceptibility phenotypes to genome assemblies, so neither could have carried out the adjustment and neither can be re-tested as it stands. The adjustment requires a sequenced genome per isolate, which is why it was available to us and not to them. The datasets large enough to make surveillance CS inference attractive are largely those in which its principal alternative explanation cannot be evaluated.

## Data availability

Raw AMR records are publicly available from the BV-BRC database at https://www.bv-brc.org (genome_amr endpoint, accessed 2026-07-31). Analysis code is available from the corresponding author on request and will be deposited in a public repository upon acceptance.

## Ethics statement

This study is based entirely on de-identified clinical isolate records from the BV-BRC public database (https://www.bv-brc.org), a US federal resource containing no personally identifiable information. No human subjects research was conducted, and no ethics approval was required.

## Acknowledgements

Analysis was performed using BV-BRC public data, NetworkX, SciPy, and NumPy.

