## Supplementary tables 1-4 for "Collateral Sensitivity Strongly Connected Components in Real-World Clinical Surveillance Data Are Confounded by Clonal Lineage"

John Goodman

OceanSparx Pty Ltd, Sydney, Australia

August 2026

**Table 1: Supplementary Table S1.** Antibiotics retained per species. The curated target list comprised 31 agents; an antibiotic was retained for a species if tested in ≥50 isolates of that species after removal of intermediate phenotypes. Checkmarks indicate retention.

| Class | Antibiotic | *K.p.* | *E.c.* | *S.a.* | *P.a.* |
|---|---|---|---|---|---|
| Carbapenems | imipenem | • | • | – | • |
| | meropenem | • | • | – | • |
| | ertapenem | • | • | – | – |
| | doripenem | • | – | – | • |
| Cephalosporins | ceftriaxone | • | • | – | – |
| | cefotaxime | • | • | – | – |
| | ceftazidime | • | • | – | • |
| | cefepime | • | • | – | • |
| Fluoroquinolones | ciprofloxacin | • | • | • | • |
| | levofloxacin | • | • | • | • |
| | moxifloxacin | – | – | • | – |
| Aminoglycosides | gentamicin | • | • | • | • |
| | amikacin | • | • | – | • |
| | tobramycin | • | • | • | • |
| $\beta$-lactam/inhibitor | piperacillin/tazobactam | • | • | – | • |
| | ampicillin/sulbactam | • | • | – | – |
| | amoxicillin/clavulanic acid | • | • | – | – |
| Polymyxins | colistin | • | • | – | • |
| | polymyxin b | • | – | – | • |
| Tetracyclines | tetracycline | • | • | • | • |
| | doxycycline | • | • | • | • |
| | minocycline | – | • | – | – |
| | tigecycline | • | • | • | – |
| Sulfonamide/TMP | trimethoprim/sulfamethoxazole | • | • | • | • |
| Glycopeptides | vancomycin | – | – | • | – |
| | teicoplanin | – | – | • | – |
| Oxazolidinones | linezolid | – | – | • | – |
| Rifamycins | rifampicin | – | – | – | – |
| Fosfomycin | fosfomycin | • | • | • | – |
| Nitrofurans | nitrofurantoin | • | • | – | – |
| Amphenicols | chloramphenicol | • | • | • | • |
| **Retained** | | **25** | **24** | **14** | **17** |
| **Isolates** | | **4,286** | **6,720** | **3,411** | **1,154** |

### Supplementary Table S2. Lineage stratification of the carbapenem–tetracycline edges

---

Contingency tables were rebuilt within each MLST sequence type using the same construction as the main analysis and pooled by Cochran–Mantel–Haenszel. Strata require $n \geq 10$ co-tested isolates. Crude values reproduce the main text to within rounding.

| Edge | Crude OR | Within-lineage OR (95% CI) | CMH $p$ | Breslow–Day $p$ |
|---|---|---|---|---|
| imipenem → tetracycline | 1.808 | 0.932 [0.632, 1.374] | 0.725 | 0.302 |
| meropenem → tetracycline | 1.819 | 0.952 [0.647, 1.400] | 0.804 | 0.211 |

Stability across the stratum-size threshold (imipenem → tetracycline):

| Minimum stratum $n$ | Strata | Isolates pooled | Within-lineage OR |
|---|---|---|---|
| 5 | 22 | 711 | 0.879 [0.601, 1.286] |
| 10 | 11 | 645 | 0.932 [0.632, 1.374] |
| 20 | 5 | 571 | 0.927 [0.622, 1.383] |

### Supplementary Table S3. Permutation control

---

Sequence-type labels were permuted across genomes 300 times, preserving the stratum-size distribution exactly, and the pooled odds ratio recomputed. This separates confounding by lineage from attenuation caused by stratification itself.

| Partition | Pooled OR |
|---|---|
| None (crude) | 1.808 |
| Size-matched random partitions, median of 300 | 1.764 |
| 95% range | [1.509, 2.131] |
| True sequence-type partition | **0.932** |

No permutation reached the observed value ($p < 0.004$, 0 of 300).

### Supplementary Table S4. Validity of the lineage assignments

---

**(a) Carbapenem resistance by sequence type**, from the same susceptibility data as the main analysis. Known carbapenemase-associated lineages are recovered without being sought.

| ST | *n* | Meropenem-resistant | |
|---|---|---|---|
| ST512 | 62 | 100.0% | single-locus variant of ST258 |
| ST147 | 94 | 77.7% | known carbapenemase clone |
| ST258 | 391 | 72.4% | globally dominant KPC clone |
| ST11 | 78 | 66.7% | dominant CR clone, Asia |
| ST37 | 63 | 9.5% | not a carbapenem-resistant lineage |
| ST45 | 46 | 4.3% | not a carbapenem-resistant lineage |
| All isolates | 2,828 | 36.4% | |

**(b) *De novo* re-typing.** Contigs were retrieved for 34 isolates, weighted to ST258 and ST307 because those strata carry 473 of the 645 co-tested isolates, and typed against the Institut Pasteur *Klebsiella* MLST scheme. Concordance with the assignments used: **32 of 33** typable genomes (97.0%); one disagreement (ST307 vs ST9830) and one genome recovering 6 of 7 loci.

**(c) Tolerance to mis-assignment.** Sequence-type labels were corrupted at random and the analysis repeated (60 replicates per level).

| Labels corrupted | Pooled OR |
|---|---|
| 0% (as measured) | 0.932 |
| 3% (observed error rate) | 0.974 |
| 10% | 1.106 |
| 25% | 1.316 |
| 50% | 1.587 |
| 100% | 1.757 |

At the observed error rate the estimate is indistinguishable from the clean one; recovering the crude value requires near-total corruption.
